# Using Deep Learning to Decipher the Impact of Telomerase Promoter Mutations on the Morpholome

**DOI:** 10.1101/2023.11.20.567914

**Authors:** Andres J. Nevarez, Anusorn Mudla, Sabrina A. Diaz, Nan Hao

## Abstract

Melanoma showcases a complex interplay of genetic alterations and cellular morphological changes during metastatic transformation. While pivotal, the role of specific mutations in dictating these changes still needs to be fully elucidated. Telomerase promoter mutations (TPMs) significantly influence melanoma’s progression, invasiveness, and resistance to various emerging treatments, including chemical inhibitors, telomerase inhibitors, targeted therapy, and immunotherapies. We aim to understand the morphological and phenotypic implications of the two dominant monoallelic TPMs, C228T and C250T, enriched in melanoma metastasis. We developed isogenic clonal cell lines containing the TPMs and utilized dual-color expression reporters steered by the endogenous Telomerase promoter, giving us allelic resolution. This approach allowed us to monitor morpholomic variations induced by these mutations. TPM-bearing cells exhibited significant morpholome differences from their wild-type counterparts, with increased allele expression patterns, augmented wound-healing rates, and unique spatiotemporal dynamics. Notably, the C250T mutation exerted more pronounced changes in the morpholome than C228T, suggesting a differential role in metastatic potential. Our findings underscore the distinct influence of TPMs on melanoma’s cellular architecture and behavior. The C250T mutation may offer a unique morpholomic and systems-driven advantage for metastasis. These insights provide a foundational understanding of how a non-coding mutation in melanoma metastasis affects the system, manifesting in cellular morpholome.

## Introduction

In the complex world of cellular biology, the interplay between genetic modifications and cellular function is a central theme. The impact of genomic alterations on cellular functions and behavior is undeniably pivotal in the onset and progression of numerous diseases. Genomic alterations, from subtle single nucleotide polymorphisms to extensive chromosomal variations, fundamentally alter cellular function, influencing a cell’s ability to survive, replicate, and interact within its environment. Understanding these alterations and their impact on the cellular morpholome is essential for developing novel image-based detection methods, therapeutic strategies, and treatment interventions. Amidst the vast genomic landscape, the promoter region of the telomerase reverse transcriptase (TERT) gene emerges as a significant focal point in melanoma. TERT promoter mutations (TPMs) have been ubiquitously identified across many human cancers, highlighting their critical role in disease progression. In melanoma, these mutations are not mere passengers but active drivers, contributing to melanoma cells’ aggressive behavior and metastatic potential [1–4].

Melanoma is a malignant neoplasm arising from melanocytes and is the most aggressive and deadliest form of skin cancer [5]. Despite constituting only about 1% of all skin cancers, melanoma accounts for most skin cancer-related deaths [6]. Over the past several decades, advancements in early detection and an improved understanding of melanoma’s molecular mechanisms have contributed to increased survival rates. However, melanoma remains a significant global health concern due to its high propensity for metastasis and resistance to therapies [7]. The metastatic stage of melanoma is primarily responsible for its high lethality, with a 5-year survival rate of only 22.5% for patients with distant metastases [8].

Several genetic and epigenetic alterations have been identified in melanoma that contribute to its aggressive behavior and metastatic potential [1,9,10]. One of the fundamental genetic alterations in melanoma is the presence of mutations in the promoter region of the TERT gene, TPMs – C228T and C250T [11,12]. TERT is a catalytic subunit of the telomerase enzyme, which maintains the telomere length and enables continuous cell division and immortalization [13]. While C228T is generally more frequent than C250T, their prevalence is nearly equal in melanoma compared to other cancer types [14]. TPMs have been found to occur early in melanoma development and become enriched in metastatic tumor sites, suggesting a functional role in the metastatic progression [15].

Melanoma represents an intriguing paradigm of malignancy in cellular morphology and systems biology, where cellular form and function shift dramatically during the metastatic stage. Metastatic melanoma cells undergo a complex and highly regulated process, including local invasion, intravasation, survival in the circulation, extravasation, and colonization at distant sites [16]. While melanoma’s genetic landscape has been extensively explored, understanding how specific genetic mutations influence the cellular architecture and networked systems is essential. The morpholomic implications of these mutations in melanoma metastasis remain poorly understood, and their potential contribution to metastatic phenotypes warrants further investigation.

This hints at TPMs’ role in reshaping cellular morpholome, which may be conducive to metastasis. Using quantitative imaging to describe metastatic cellular properties [17], we define the metastatic morpholome as the complete set of single-cell morphological features encompassing dynamic processes such as motility and static measurements such as intensity or deep learned latent features. Recent studies have begun to elucidate the impact of TPMs on melanoma progression, highlighting their influence on various cellular processes, including cell proliferation, migration, and invasion [18,19]. Despite these advancements, a significant limitation lies in the reliance on cancer cell lines or patient-derived tumor samples, which inherently contain many background mutations. These confounding factors potentially obscure the precise effects of TPMs on metastatic phenotypes, necessitating a more isolated and controlled examination.

In response, we meticulously engineer isogenic clonal cell lines with monoallelic TPMs, eliminating the influence of background mutations. This approach allows for a more refined observation of the morpholome implications of these mutations, providing a clearer insight into their role in metastatic behavior. This innovative technique facilitates a comprehensive exploration of the functional clonal consequences of C228T and C250T mutations, offering a detailed understanding of the induced morpholomic variations.

Our findings reveal pronounced morpholomic shifts between cells harboring TPM variants and their wild-type counterparts. These morpholomic variations encompass diverse aspects, from allele expression patterns to morphological changes, enhanced wound-healing rates, and distinctive spatiotemporal dynamics. A notable observation is the distinct divergence in the influence of C250T and C228T mutations on metastatic phenotypes, countering the previously held belief of their functional equivalence. The C250T mutation demonstrates significant morpholomic shifts, potentially amplifying its metastatic capabilities.

Embracing a symbolic perspective, this work likens the introduction of a TPM to a rock cast into a stream. This seemingly minor mutation sends ripples throughout the cellular morpholome, altering the trajectory and interactions of cellular behavior. This study’s design, centered on engineering cells with a non-coding promoter mutation, enables a focused observation and analysis of the mutation’s effects on the cellular morpholome. This approach unveils the multifaceted implications of TPMs in melanoma progression and metastasis, offering a novel and enriched perspective on the intricate interplay of genetic modifications and the morpholome.

## Results

### C250T increases metastatic potential and penetrance in multiple organs

A clonal composition change in cancer progression from primary to metastatic disease enriches cells harboring the TPMs [5]. This clonal evolution suggests a functional advantage for metastatic potential and penetrance due to TPM status. To quantify metastatic potential and penetrance of TPMs *in vivo*, we mined the large-scale pan-cancer study MetMap [20]. MetMap does not include the TPM statuses of their cell lines; instead, we searched cell databases and primary literature to identify the TPM statuses of cell lines. We acknowledge that the size of our C228T and C250T samples used in the subsequent analysis is far from the 500 human cancer cell lines used in the MetMap study because we are limited by the fact that TPM status can only be found using whole genome or Sanger sequencing. Clinics and research groups often use exome sequencing for cancer mutation profiling. We could not stratify C250T and C228T by cancer type due to a lack of TPM information for many of the 500 cell lines used in the MetMap study. However, we did find that C250T had higher penetrance and potential than the C228T pan-cancer in all organs (**Fig. 1, A, and G**). Breaking down metastatic potential and penetrance by each organ for TPMs, we see that C250T has significantly higher penetrance in the lungs, liver, and brain (**Fig. 1, B, C, and D**). At the same time, the kidney and bone show no difference (**Fig. 1, E, and F**). In addition, C250T has significantly higher metastatic potential in the lung and liver compared to the kidney, bone, and brain, which show no difference (**Fig. 1, H-L**). We found that TPMs do not correlate with aneuploidy, mutational burden, or replication rate of the samples used from the MetMap data. It should be noted that the difference in the metastatic potential of TPMs in melanoma was corroborated by a recent study using interpretable deep-learned models to identify high and low-efficiency metastatic melanoma single-cell properties [21]. The interpretable deep-learned model predicted A375, which harbors the C250T mutation [22], to have increased metastatic potential in a patient-derived mouse model. This was confirmed in the metastatic mouse model established by the landmark study of metastatic melanoma [23]. While the consensus is that TPMs, C250T, and C228T are genetically identical, these results suggest they differ *in vivo*. Furthermore, since C250T occurs in higher frequency in melanoma, a cancer type known for its aggressive metastatic stage [24], it may significantly impact metastatic phenotypes. Our meta-analysis shows that pan-cancer tumors harboring the C250T TPM have higher metastatic potential and penetrance. However, we are not extending this to claim that C250T is solely causative in increasing metastatic potential and penetrance over C228T.

**Figure 1.**
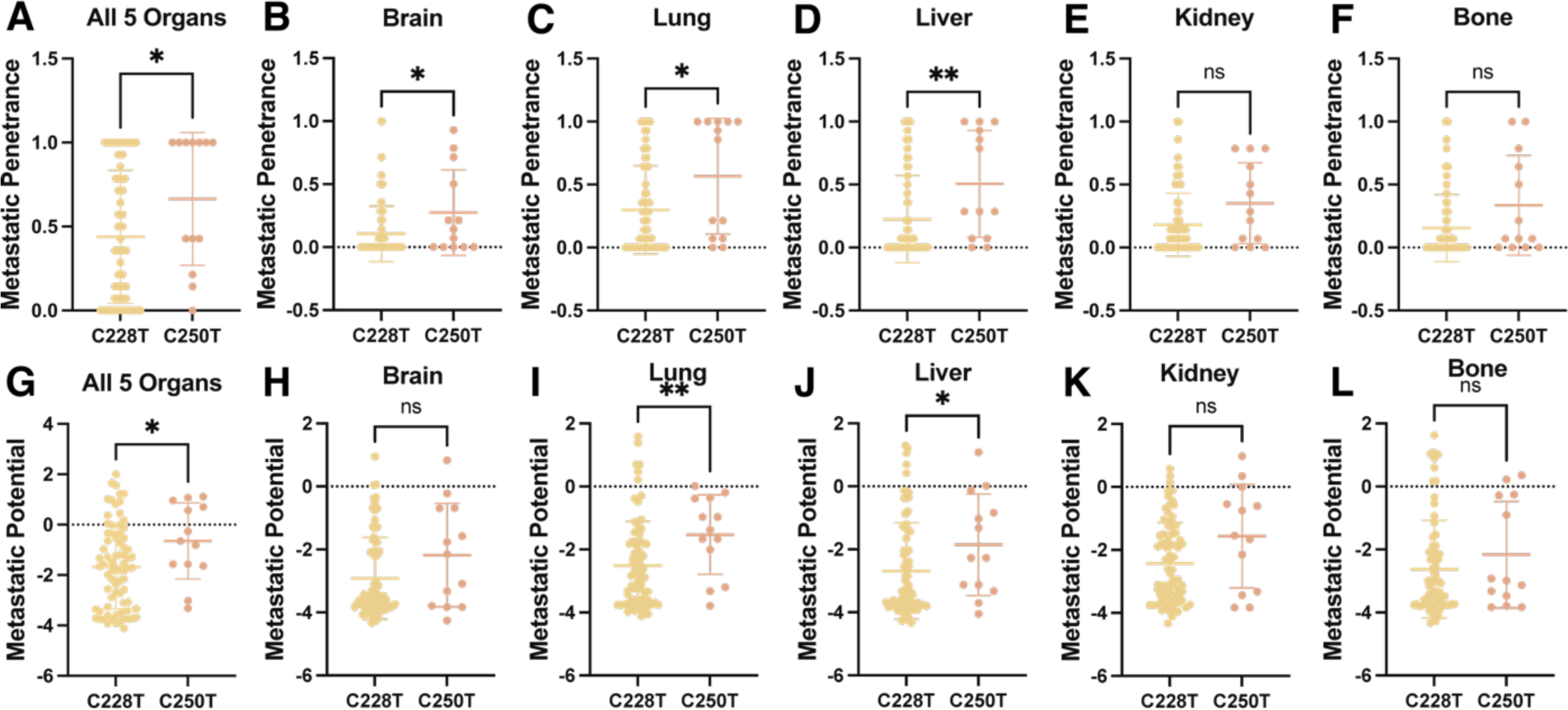
Analysis of mined information from MetMap database for quantified metastatic potential and penetrance of cell lines harboring Telomerase Promoter Mutations stratified by C228T and C250T. C228T n = 76, and C250T n = 13. **(A)** Metastatic Penetrance comparison of C228T and C250T aggregated by all organs used in the MetMap database, p =0.0330. **(B, C, D, E, F)** Metastatic Penetrance comparison of C228T and C250T separated by organ Brain p = 0.0240, Lung p = 0.0326, Liver p =0.0055, Kidney p = 0.0615, and Bone p = 0.0766. **(G)** Metastatic Potential comparison of C228T and C250T aggregated by all organs used in the MetMap database, p = 0.0326. **(H, I, J, K, L)** Metastatic Potential comparison of C228T and C250T separated by organ Brain p = 0.1555, Lung p = 0.0099, Liver p = 0.0347, Kidney p = 0.0819, and Bone p = 0.3706. All p values from the Mann-Whitney test.

### TPMs increased allele expression, manifesting in multiple changes in cellular properties

To dissect the effect of each TPM on metastatically relevant cellular properties, we introduced these mutations in HEK 293T cells, which have a minimal mutational burden and unlimited replicative ability due to viral integration rather than TERT mutation or amplification [25]. Since TPMs are mutually exclusive and occur only on one allele, we made individual mutations but not double mutations [12,26]. Importantly, we designed this approach to eliminate potential heterogeneity due to genetics, such as in primary cancer cells or metastatic melanoma cell lines, which would confound our analysis. To this end, HEK 293T *TERT*^WT/WT^, *TERT*^C228T/WT^, and *TERT*^C250T/WT^ single-cell clones were engineered with spectrally distinct fluorescent expression reporters on each allele under the endogenous promoter and at the endogenous loci **(Fig. 2, A and B, and Supp Fig. 1)**. This was meticulously done such that the endogenous TERT promoter controlled the reporters. Furthermore, to preserve the natural genomic context and ensure the native regulatory mechanisms remained intact, these fluorescent reporters were incorporated precisely at the endogenous loci of the TERT gene in the genome; we elected to use P2A [27] so that the expression of the endogenous Telomerase open reading frame is left intact. This setup allowed for an accurate representation of the impact of the respective mutations on gene expression, considering the native chromosomal environment and its potential epigenetic effects. Using quantitative fluorescence microscopy, we observed that the mean expression levels from the TPM allele increased by ∼1.4x over the corresponding WT allele **(Fig. 2C)**, consistent with previous studies [26,28–30]. We also observed a significant increase in the mean expression of the C250T over the C228T TPM allele, which has yet to be reported. We quantified the impact of TPMs on the cell-to-cell variation of allele expression using the Fano Factor **(Fig. 2D)** [31] seen in the single-cell scatter plot **(Fig. 2E)**. While both TPMs increased the variations in gene expression relative to WT, and C250T showed a more dramatic effect than C228T.

**Figure 2.**
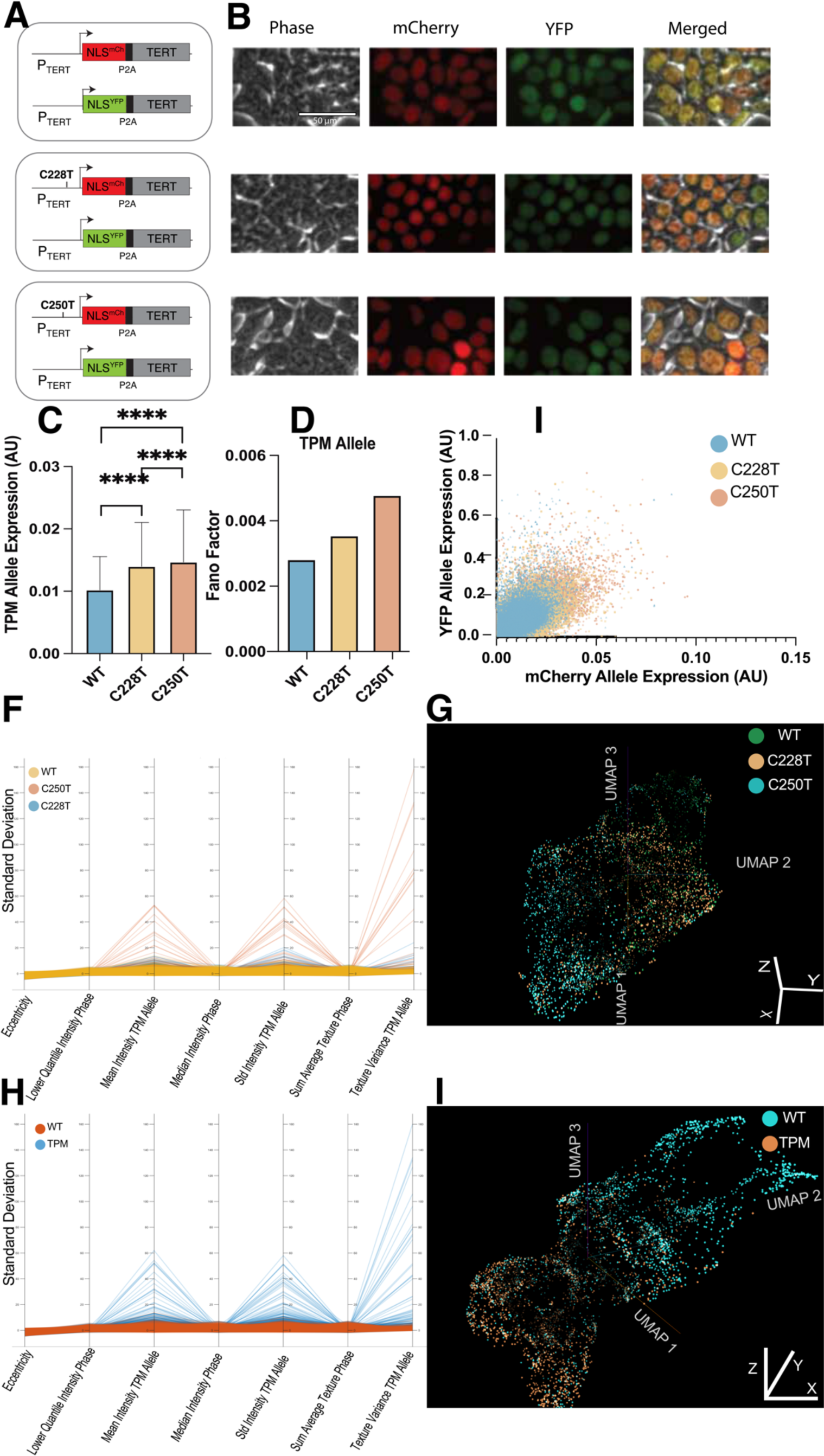
Introduction of the TPMs illicit a response in the morpholome. **(A)** Schematic of the engineered genomic structure of the cell lines. **(B)** False-colored image montage of representative cells for Telomerase promoter mutants, C228T, C250T, and WT cell lines. Telomerase gene expression separated by allele for n = 67,000 cells. **(C)** Bar graph showing the difference in mean mCherry Allele expression of the mutant allele containing control WT allele and the promoter mutation C228T and C250T, respectively. WT displayed significantly lower expression than C228T p <0.0001 and C250T <0.0001. While C228T showed significantly lower expression than C250T <0.0001 (unpaired t-tests). **(D)** Bar graph showing the allele expression variance for the mCherry Allele quantified using the Fano Factor calculation. **(E)** Scatter plot overlaying the WT, C228T, and C250T allele expression. Each data point represents a single cell. **(F)** The standard deviation of parallel coordinates of the top 5 discriminative features from a Random Forest trained on individual mutation status features. **(G)** 3D UMAP of the Average Pool 4 latent embeddings in a modified Resnet-50 model trained on phase contrast images labeled by individual mutation status. **(H)** Standard deviation parallel coordinates of the top 5 discriminative features from a Random Forest trained on TPM (C228T and C250T) and WT features. **(I)** 3D UMAP of the Average Pool 4 latent embeddings in a modified Resnet-50 model trained on phase contrast images labeled by TPM (C228T and C250T) and WT.

Building from the unexpected allele expression heterogeneity, we sought to uncover differences in multiple cellular features that define the morpholome, such as intensity, shape, and texture of the cell and nucleus, a total of 523 engineered cell features **(Supp Fig. 2)**. We used the CellProfiler Cellpose plug-in to segment the entire cell **(Supp Fig. 3)** using the phase contrast channel using the Cyto2 model [32,33] to quantify cellular features and a nuclear reporter to get nuclear features. We trained a Random Forest classifier model to discriminate between WT, C228T, and C250T cells using the 523 morpholome features. We found the top features of the Random Forest model related to the expression of the TPM allele and intensity and texture in the phase contrast image. We plotted the parallel coordinates of the standard deviation of the top 6 normalized features [34] as individual mutation status classes (C228T, C250T, and WT) **(Fig. 2F)** with an accuracy of 67.44% **(Supp Fig. 4A)** and grouping the C228T and C250T mutational status cell lines into a TPM class **(Fig. 2H)** with an accuracy of 78% **(Supp Fig. 4B)**.

Following up on the features related to phase contrast images, we trained modified Resnet-50 models to discriminate between WT, C228T, and C250T and grouping the C228T and C250T mutational status cell lines into a TPM class [35]. We extracted the latent embeddings of the penultimate layer from the models and used the UMAP dimensionality reduction [36] to project these features in three dimensions as individual mutation status classes (C228T, C250T, and WT) **(Fig. 2G)** and grouped the C228T and C250T mutational status cell lines into a TPM class **(Fig. 2I and Supp Fig. 5)**. The TPM vs WT model achieved significantly better accuracy than the individual mutation status classes.

### Defining the impact of TPMs on the morpholome

We sought a different experimental modality to highlight cell appearance changes in a non-2D cell culture environment. Imaging Flow Cytometry (IFC) allows the cells to be suspended in solution, experiencing different mechanical properties, which increases the Epithelial to mesenchymal transition (EMT) genes [37]. The increased expression in these pathways may illicit a more pronounced response in cell morphology. The pairing of deep learning and machine learning with IFC has been used to interrogate the metastatic stage in primary samples as part of the pathology pipeline. IFC can probe the cell using cell light scatter and brightfield images. Both require no labels and have been used for cancer classification with deep and machine-learning [35,38–40].

Here, we leverage the power of the ImageStream imaging flow cytometry platform [41] and Deepometry [35] to identify differences between WT cells and cells harboring the TPMs and further subtle differences between cells harboring C228T and C250T. We found that we can discriminate between the three cell lines using label-free images alone **(Fig. 3A)**. We first examined the latent embeddings of the penultimate layer before the classification grouping the cell lines as TPM harboring cells vs. WT cells reducing the dimensions using 3D UMAP **(Fig. 3B and Supp Fig. 6A)** and 3D PHATE [42] **(Fig. 3C and Supp Fig. 6B)** plots for trained model has an overall accuracy of 94.4%. We observe two clusters of cells in the UMAP plot, showcasing that the TPM causes system-wide changes manifesting in the brightfield and side-scatter images. The PHATE reduction of the exact latent embeddings allows us to visualize transition morphological states from WT to TPM. Next, we trained a model grouping the mutation statuses individually with an accuracy of 66.5%. We still observe two distinct clusters in the 3D UMAP **(Fig. 3D and Supp Fig. 7A)**; however, C250T and C228T do not cluster individually within the supercluster. We visualize the latent features using PHATE **(Fig. 3E and Supp Fig. 7B)** and observe the transition states in granular detail. Similar to the UMAP, there are two clusters of cells, with one cluster predominantly related to WT, while the other corresponds to the C228T and C250T features. However, in the transition zone between the two clusters, we observe WT and C250T dominating this zone.

**Figure 3.**
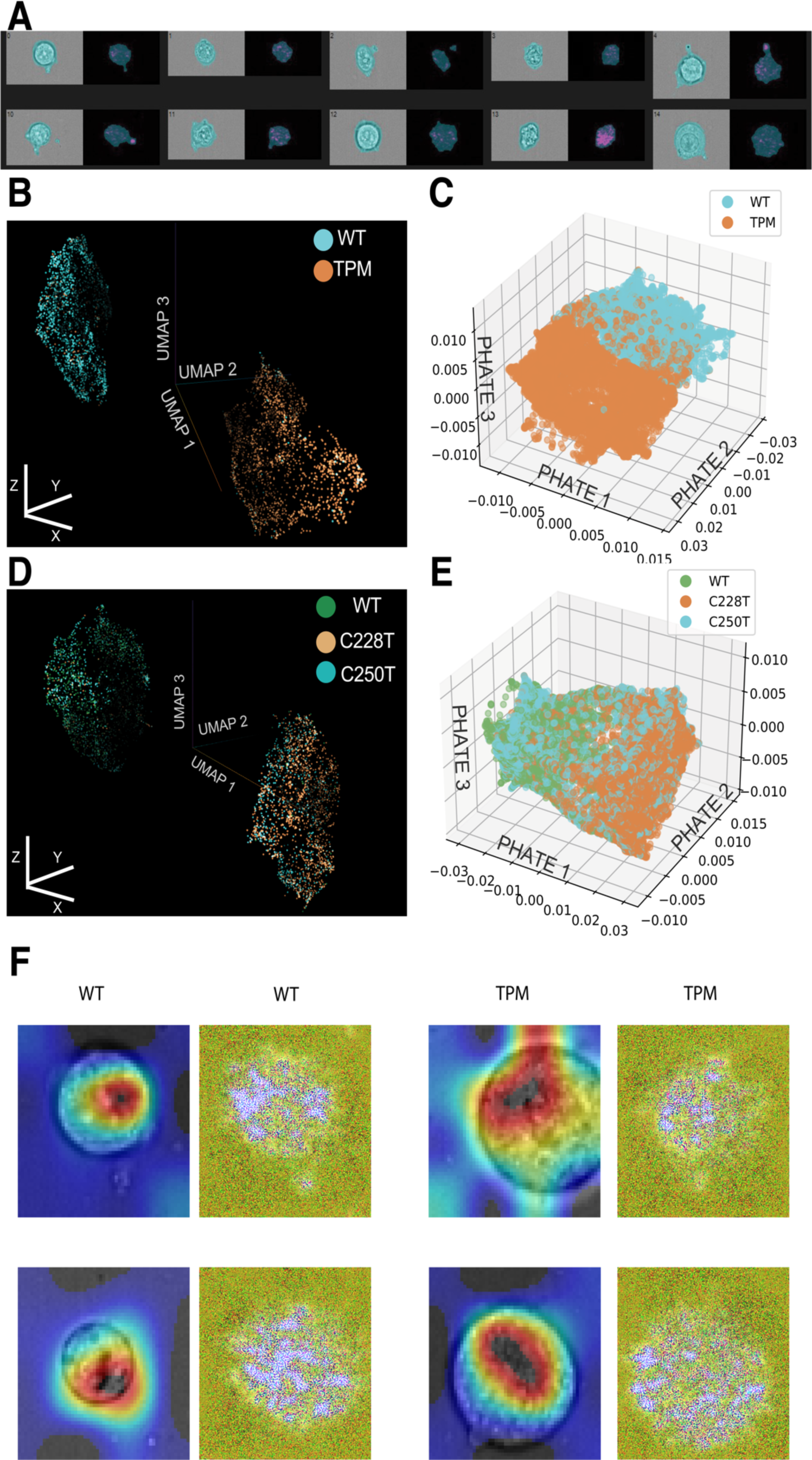
The latent features from the deep-learned features of label-free images allow us to discriminate between the cell lines and visualize the morphological transition states. **(A)** Representative image montage of cells using the IFC; Brightfield is in gray, the Side-scatter image is falsely colored in magenta, and the cell mask is overlayed in blue. **(B)** 3D UMAP **(C)** 3D PHATE of the latent embeddings of the Average Pool 4 in a modified Resnet-50 model trained on label-free images labeled by TPM (C228T and C250T) and WT. **(D)** 3D UMAP **(E)** 3D PHATE of the latent embeddings of the Average Pool 4 in a modified Resnet-50 model trained on label-free images labeled by individual mutation status. **(F)** Representative images of Grad-Cam overlayed (heatmap) on the brightfield (Grey) image for the final convolution layer and corresponding Deepdreamed images in yellow.

Visualizing the Gradient-weighted Class Activation Mapping (Grad-Cam) [43], or what the AI model sees as most relevant and essential when predicting an image class (i.e., TPM vs WT), homes in subtle yet significant differences observed in the brightfield images **(Fig. 3F)**. For both WT and TPM, we see the focus of the Grad-Cam on the interior for highly textured and granular features. For the TPM cells, we see the Grad-Cam has a broader area in the cell, suggesting larger texture and granularity differences than the WT cells, which have a smaller focus on the interior parts of the cell. To further our understanding of the morpholomic features of our model, we then visualized the Deepdreamed images [44], which visualizes the patterns our convolutional neural net learned. We see, again, the TPM features related to texture and granularity **(Fig. 3F)**. There is a more focal point in the TPMs vs WT. In contrast, WT in the Deepdreamed images follows the teal in the Grad-Cam, whereas the TPMs are more specific, focusing on points of interest spatially in the interior of the cell. These two techniques allow us to delineate the impact of the TPMs on the morpholome. Additionally, we trained a Random Forest Classifier on the quantified engineered cell features; we achieved an accuracy of 94% **(Supp Fig. 8A)** with the top features relating to brightfield image morphology, particularly texture, intracellular light scattering (intensity), and sub-cellular spatial distribution differences (radial distribution of Zernike features) **(Supp Fig. 8B)**. Still, size was not different between TPM and WT. Combining these three techniques, we can confidently say the differences in morpholomic profiles between TPM and WT cells are related to changes in texture and intensity features in the brightfield and side-scatter images.

### TPMs increase wound healing spatiotemporal migration dynamics

Following up on our findings from Figure 1 regarding C250T increasing penetrance *in vivo*, we wanted to investigate if C250T increases collective cell migration using the wound healing assay. Wounds are a breach in cells that are usually linked tightly to each other to form a protective barrier and are suddenly separated. Similarly, cells detach from their neighbors in metastasis and adopt a migratory behavior to reach new locations. The wound repair program is suspected of equipping both types of cells to survive this anchorless state [45]. This allows cells to move into the breach and make new tissues, enabling metastatic cells to detach and colonize new destinations. Increased Telomerase expression has been shown to promote the epithelial to mesenchymal transition and three-dimensional invasion through Matrigel *in vitro* [46,47]. The close connection of Telomerase and WNT/μ-Catenin pathway is connected to invasiveness and stemness in cancer [48–50].

We investigated spatiotemporal dynamics of collective cell migration between the three clonal cell lines via *in vitro* wound healing using previously published software [51]. The WT cell line had a significantly lower healing rate than C228T and C250T. C250T showed a substantially higher healing rate than C228T **(Fig. 4A)**. The overall distances migrated showed similar patterns, with WT having significantly lower spaces than C228T and C250T. In contrast, C250T migrated further than C228T **(Fig. 4B)**.

**Figure 4.**
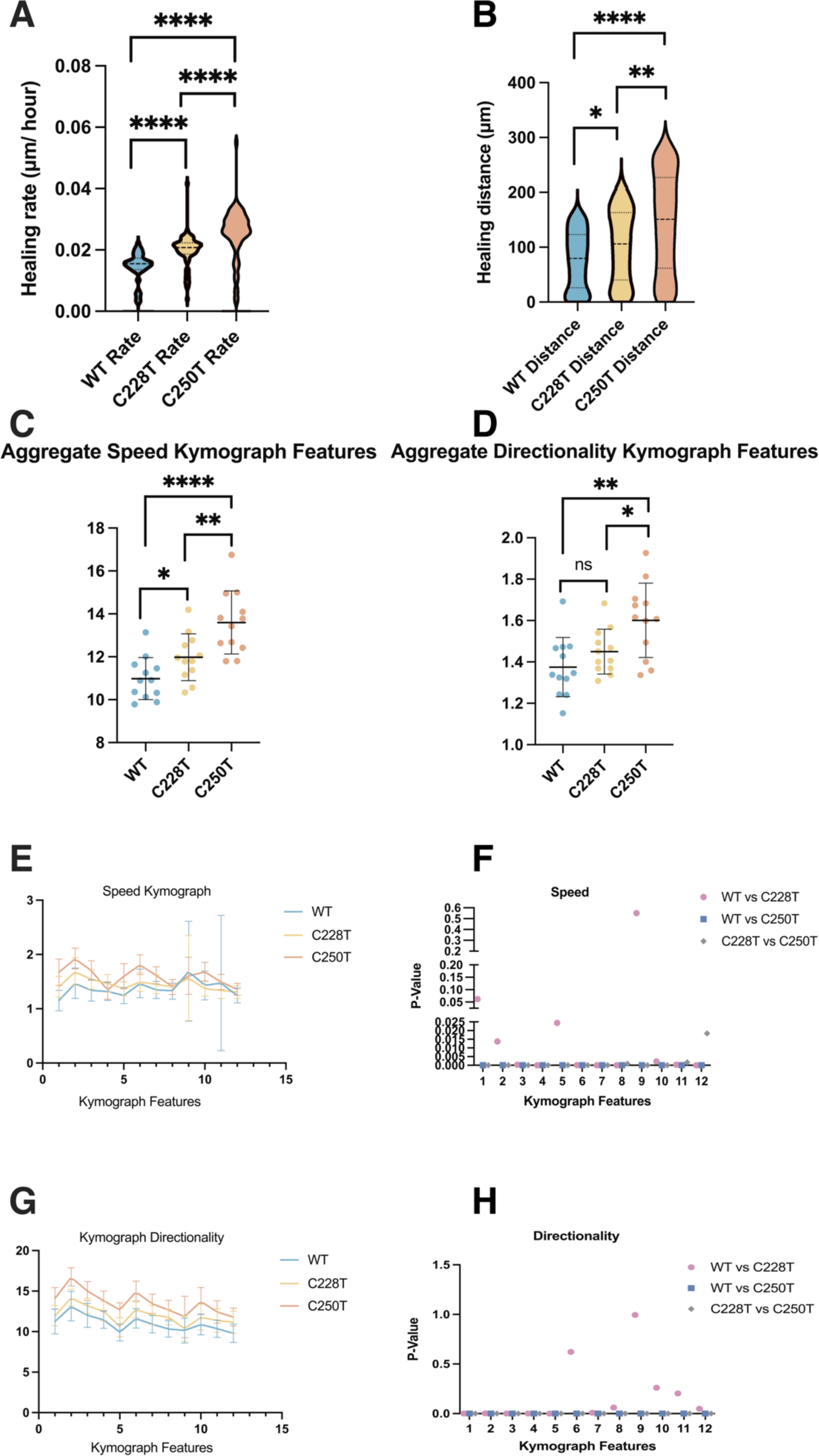
C250T has increased spatiotemporal monolayer migration using live-cell *in vitro* wound-healing assay. n = 38 movies for each cell line. **(A)** Violin plot comparison of healing rates across cell lines. WT compared to C228T p <0.0001, WT compared to C250T p <0.0001, and C250T compared to C228T p <0.0001. **(B)** Total distance in microns is the leading edge covered throughout the movies. WT compared to C228T p = 0.0175, WT compared to C250T p = 0.0047, and C250T compared to C228T p <0.0001. Aggregate comparisons of the average of each of the 12 kymograph features for all movies across cell lines. **(C)** Speed of each of the 12 averaged figures. WT compared to C228T p = 0.1600, WT compared to C250T p = 0.0036, and C250T compared to C228T p = 0.0332. **(D)** Directionality of each of the 12 averaged figures. WT compared to C228T p = 0.0242, WT compared to C250T p <0.0001, and C250T compared to C228T p = 0.0083. **(E)** Graph showing the speed of individual 12 kymographs features averaged among the 38 movies with SD error bars. **(F)** Graph depicting p-values from multiple Mann-Whitney tests for speed features. WT compared to C228T were significantly different in features 2-8 and 10-12 with p values 0.013734, 0.000344, 0.000010, 0.024413, 0.000085, 0.000010, <0.000001 and 0.002214, 0.000359, and 0.000014 respectively. Features 1 and 9 were not significantly different, with p-values of 0.061604 and 0.550616, respectively. WT compared to C250T were significantly different in all 12 features, p <0.000001. C228T compared to C250T differed significantly in all features with corresponding p values of <0.000001, <0.000001, <0.000001, 0.000012, <0.000001, <0.000001, 0.000006, 0.000887, 0.000008, 0.000003, 0.001708, 0.018402. **(G)** Graph showing the directionality of individual 12 kymographs features averaged among the 38 movies with SD error bars. **(H)** WT compared to C228T differed significantly in directionality features 1-5 and 7 with p values <0.000001, 0.001646, 0.000027, 0.000670, 0.000093, and 0.008123 respectively. While features 6 and 8-12 were not significantly different with corresponding p values of 0.621746, 0.060135, 0.993789, 0.260206, 0.203475, and 0.048283. WT compared to C250T were significantly different in all 12 features, p <0.000001. C228T compared to C250T differed significantly in all 12 features p <0.000001. All p values from Mann-Whitney tests.

We also examined the averaged speed and directionality of the kymographs across experiments. We found that C228T showed a significantly higher speed than WT but a very modest difference in directionality compared to WT. C250T, however, showed substantially higher speed and directionality than either WT or C228T **(Fig. 4, C and D)**. We then investigated the speed in each of the 12 features of the kymographs individually. We found that WT and C228T differed in only six features, whereas C250T showed significant differences in all 12 features compared to WT or C228T **(Fig. 4, E, and F)**. We next examined the speed of each of the 12 features. C228T displayed significant differences from WT in 10/12 features, whereas C250T differed from WT or C228T in all 12 features **(Fig. 4, G, and H)**.

We showed that C250T has a considerable advantage in healing rate and overall distance covered in an *in vitro* wound-healing assay, followed by C228T and WT. Furthermore, C250T has notable spatiotemporal differences in speed and directionality propagating from the wound edge into the monolayer compared to WT and C228T. C228T lags behind C250T in all metrics but is still significantly faster and migrated further than WT, with significant spatiotemporal differences in nearly all 12 features. These results align with the differences observed in TERT expression from C228T and C250T **(Fig. 2)**, which may underlie the higher metastatic potential and penetrance of C250T compared to C228T **(Fig. 1)**.

## Discussion

While the reactivation of Telomerase is crucial for melanoma development, there is no direct link between long telomeres and metastatic potential [67] in melanoma. This suggests that, besides enabling replicative immortality, Telomerase may have alternative functions [68] relevant to withstanding cellular stress during metastasis. TPMs are enriched in metastatic tumors across various cancer types, signifying a stereotypical advantage in surviving cellular stress for disseminating cells. Although both C228T and C250T TPMs generate a new *de novo* Erythroblast Transformation Specific (ETS) transcriptional binding site, their prevalence differs across cancers, with C228T being predominant, except in melanoma [18,19]. Metastatic melanoma exhibits high metastatic potential and is linked to changes in morphological cellular properties [21]. However, the specific metastatically relevant cellular morpholome properties affected by TPMs remain undefined.

This study uses live cell imaging, machine learning, and deep learning to investigate how TPMs influence metastatically relevant cellular morpholome properties. We first exploited the extensive database of quantified *in vivo* metastatic fitness in MetMap, revealing through meta-analysis that tumors with C250T TPM have increased metastatic potential and penetrance across all studied organs than those with C228T TPM **(Fig. 1)**. While the two TPMs are regarded as genetically identical, their differential impacts *in vivo* suggest an underlying divergence in function. Notably, the heightened prevalence of the C250T mutation in aggressive metastatic melanomas underscores the significant clinical implications of this mutation in tumor progression and metastasis. Having established *in vivo* differences, we sought to observe the morpholome differences.

To understand the effect of TPMs on the morpholome without the confounding influence of mutations and copy number variance, we engineered isogenic cell lines containing C228T or C250T TPMs at the endogenous locus under the endogenous promoter to examine the impact of TPMs *in vitro*. C250T confers increased expression and variance of TERT expression than C228T. C250T has higher levels of variance in the top discriminative features of our Random Forest model than C228T. Stochastic gene expression allows clonal populations to exhibit phenotypic heterogeneity, possibly aiding in metastatic dissemination [52–55]. We posit that the increase in the heterogeneity of TERT expression could be a barrier to effective therapy [56], which might account for the metastatic cell’s ability to survive multiple stressors in metastatic dissemination. Moreover, using machine-learned models based on over 500 cell features, we could distinguish between C250T and C228T with high accuracy and low false positives **(Fig 2)**. Interpreting the top discriminative features of the model further elucidated the morpholomic differences induced by these TPMs. TPM’s morpholome effect is especially striking in the deep-learned model trained on label-free phase contrast images, where we can discriminate more accurately.

To further understand cell morphology in a non-2D culture, we used a different live cell imaging modality, Imaging Flow Cytometry (IFC). Using the IFC modality, we have delineated morphological differences between these isogenic cell lines differing in only one base pair. Using label-free images to define the morpholome alone was sufficient to distinguish these cell types, suggesting profound morphological changes induced by the TPMs **(Fig. 3)**. Nonetheless, discerning between the two TPMs, C228T and C250T, remains challenging, indicating the subtle differences between these mutations. However, we visualized the discriminatory feature patterns of the deep-learned model using GradCam and Deepdream, allowing us to interpret what cellular properties changed: cellular texture and intracellular intensity or light-scattering properties. We can accurately discriminate between TPM and WT cells, meaning the latent embeddings of TPMs are stereotypical image features of metastasis. While C228T is more common in patients, the C250T has the plasticity to transition back and forth, supporting the increased *in vivo* fitness **(Fig. 1).** This illustrates how a non-coding mutation affects the single-cell system beyond the increase in gene expression. While there is considerable work to translate this to clinical pipelines, these results show the capability of label-free morphological features to identify cells harboring TPMs to direct treatment plans.

We sought to understand how these static morpholome profiles held up during dynamic processes, such as migration, which is critical to metastasis. Cells with the C250T mutation displayed increased migration rate, distance traveled, coordination, and speed at the leading edge and throughout the monolayer, likely mirroring their increased metastatic abilities *in vivo*. Furthermore, the spatiotemporal dynamics between the cell types followed the same trend we have seen in our other results, with C250T having the most increased migration dynamics, followed by C228T and then the WT cells. The intricate interplay between the TPMs and the wound repair machinery, especially given the role in facilitating metastatic dissemination, is intriguing. Our results hint at the potential therapeutic relevance of targeting the wound repair pathways in cancers harboring TPMs **(Fig. 4)**.

In the evolving landscape of metastatic research, understanding metastasis’s genetic and morpholomic underpinnings is pivotal. Our study sheds light on the functional impact of TPMs on morpholome cellular properties related to metastatic potential, cellular expression, and migration dynamics. Though our analysis clearly shows increased robustness in C250T-bearing cells, it is vital to note that we are not purporting that the mutation is the sole causative factor to metastatic potential, but rather, how it affects the cellular system as readout from the morpholome. Our research underscores the multifaceted impacts of TPMs, notably C250T, in promoting metastatic potential, altering gene expression dynamics, and inducing distinct cellular phenotypes. While we have charted new territories in understanding the differential *in vitro* roles of TPMs, further exploration into their mechanistic underpinnings will be instrumental in devising novel therapeutic interventions for TPM-bearing cancers. Given the ever-important proper screening of biopsy samples, we envision developing morpholome-based AI models based on imaging flow cytometers like the Amnis ImageStream [41], BD S8 with Cellview [57], and Deepcell’s REM-I system [58], to provide a metastatic potential score that associates with the increased potential derived from the TPMs.

## Methods

### Cell culture and origin

HEK293T was ordered from ATCC (ATCC CRL-11268) and tested negative for Mycoplasma using a Mycoplasma detection kit (Southern Biotech). HEK293T cells were cultured in Dulbecco minimal essential medium (DMEM: Thermo Scientific HyClone #SH30022FS) supplemented with 10% fetal bovine serum, four mM L-glutamine, 100 I.U./ml penicillin, and 100 mg/ml streptomycin at 37 °C, 5% CO2 and 90% humidity.

### Transfection

The transfections were performed with a 1 mg DNA: 2 ml Fugene HD (Promega E2311) ratio. Cells were seeded at 300,000 cells/well in a six-well plate for 18 hours before transfection. Two days after transfection, puromycin was added to the medium at 1 mg/ml, and cells were selected for two days. Survival cells were grown for another seven days before being sorted with FACS into 96-well and expanded into monoclonal cell lines. We plated 300,000 cells/well in a 6-well plate overnight, then transfected with 1.25 ug gRNA and 1.25 ug linearized donor with 5 ul FugeneHD in 100 ul final volume in Opti-MEM.

### Construct designs and homogenous clonal cell line creation

We followed the CRISPR/Cas9 protocol [59] to construct the reporter cell line. In general, the gRNAs were designed by the online CRISPR tool (http://crispr.mit.edu), and the DNA oligos were ordered from Eurofins Genomics, annealed, and cloned into pSpCas9(BB)P2A-Puro (Addgene #48139) vector plasmids. gRNA plasmids were transfected into HEK293T cells and tested for gRNA efficiency using the T7 endonuclease assay. Only the most efficient gRNA was used with the donor DNA. The donor plasmids were constructed using the Gibson assembly method. We used site-directed in-vitro mutagenesis to make a synonymous substitution in the donor plasmids to avoid gRNA recognition and Cas9 cutting of the linearized donor DNA. We first developed a nuclear marker cell line by inserting the nuclear localization signal and two copies of infrared fluorescent protein (NLS-2xiRFP) [60] under the endogenous actin promoter followed by a P2A spacer in HEK293T cells. This cell line ensured a constitutive expression without introducing an exogenous strong constitutive promoter and greatly assisted cell segmentation and tracking. Briefly, the gRNA and the linearized donor DNA were transfected into HEK 293T cells, and the transfected cells were screened with 1 mg/ml puromycin for two days. The cells were allowed to grow for an additional five days before being sorted by FACS. The fluorescently positive cells were sorted as single cells into 96-well plates [59]. We collected at least 500 single cells. We grew the cells for an additional three weeks to obtain homogenous clones. On average, about 30% of cells formed colonies, and all were screened for a fluorescent signal with the microscope. A minimum of 10 clones were then genotyped and checked for homozygosity and correct integration using at least three pairs of primers and confirmed with sequencing. Positive clones were further validated with a western blot to ensure valid protein expression. After construction and validation, the engineered single-clonal cell line was assigned a unique identification number, entered into our electronic database, and stored in liquid nitrogen with a cryoprotectant. The same procedure was performed for CRISPR-based tagging of the other allele of Telomerase. Cells were screened with puromycin and sorted by FACS to generate monoclonal cell lines.

### Flow cytometry and single-cell sorting

Cells were released via trypsinization and filtered through a cell strainer to create cell populations “cloned” from a single cell (Fischer; 07-201-430). To ensure a single-cell solution, cells were resuspended in HBH and analyzed using a benchtop BD FACSAria Fusion flow cytometer (BD Biosciences, Franklin Lakes, NJ). Single-cell clone generation of WT and TPM cell lines were sorted using fluorescent-activated cell Sorting into 96-well plates. Acquired flow cytometry data were all analyzed with FlowJo software (Tree Star). All cells were gated on FSC, SSC, and iRFP, and then mCherry and YFP took cells from the bulk of the populations, not the outliers. Single cells were identified via phase contrast and fluorescence microscopy. Once confluent within the well, the clonal populations were released via trypsinization, transferred to individual cell culture dishes, and allowed to expand until confluency.

### Live cell imaging

The imaging medium is phenol red-free DMEM (Life Technology-Gibco) with the same supplements as the regular culture medium. Live cell phase contrast and fluorescence imaging were performed on a Nikon Ti microscope with an environmental chamber held at 37oC and 5% CO2 in 20x magnification (pixel size of 0.803μm). Cells were seeded in a 24-well polystyrene dish with a cell count of 240,000 cells in 4 mL of phenol-red free DMEM per dish. The dish was then placed in a 37°C incubator for 24 hours to allow cells to recover from trypsinization before imaging. Snapshot images were taken before the time-lapse. Time-lapse imaging was performed for 3-4 days to allow for multiple cell divisions.

### Collective cell migration

Cells were grown in a 2-well culture insert (ibidi; 81176) until a cell monolayer was established using Phenol-red free DMEM. The insert was removed, and 1ml medium was added while monolayer positions were imaged every 10 minutes over 24 hours. Each time-lapse movie was inspected for quality over 24 hours. If the movie time needed to be cropped due to monolayers colliding, we removed these time frames to not affect the analysis; all movies had the same time length. Step-by-step instructions and code availability [51]. The plots were generated using PRISM 9.

### Cell segmentation and feature quantification

Background correction was performed using ImageJ’s Rolling Ball background subtraction algorithm with a 15-pixel radius [61]. We used Cellpose and CellProfiler [62] for automated cellular and nuclear segmentation using phase image and the nuclear iRFP image. We also excluded non-cell objects by size and shape selection. Cell segmentations were then subject to manual inspection, and segmented objects that did not correspond to cells were eliminated. Cells in active replication were removed based on size.

### Imaging flow cytometry acquisition

Cells were released via trypsinization to image single cells and filtered through a cell strainer (Fischer; 07-201-430). Cells were resuspended in HBH and analyzed using a benchtop ImageStream X platform to capture images of live HEK 293T cells. For each cell, we captured images of brightfield and side scatter images. After image acquisition, we used the IDEAS analysis tool (accompanying the ImageStream X software) to discard multiple cells or debris, omitting them from further analysis. ImageStream settings: Sample volume: 1 ml. Flow diameter: 7 mm. The velocity of flow: 44 m s 1. Resolution: 0.5 mm. Magnification: 40. Camera sensitivity: 256 on all channels. Camera gain: 1. Brightfield LED intensity: 88 mW. Darkfield laser intensity: 1 mW. 405 nm laser set to 15 mW.

### Quantification and statistical analysis of Metastatic potential and penetrance

The mining of cell databases and primary papers determined TPM status. Plots were made in PRISM 9^®^.

### General Image Analysis

All computations were performed on an AMD Ryzen 5 3600 CPU @ 4.1 GHz machine with an NVIDIA^®^ RTX^®^ 3060 GPU running Windows 10 Professional. This workstation was built as a moderate deep-learning workstation accessible to most research budgets.

## Acknowledgments

We thank Yishaia Zabary for allowing access to the updated Spatiotemporal monolayer migration software and providing advice about parameters. We thank Dr. Phuc Nguyen and Dr. Paula Godoy in the Hao lab for providing feedback on the manuscript. AN would like to thank his funders, the HHMI Gilliam and Ford Predoctoral Fellowship. AN would like to thank GmAP Fred Hutch Grant for funding the GPU in the workstation and quantitative courses for developing AI models. This work was supported by NIH R01 GM111458 (to N.H.) and NIH R01 GM144595 (to N. H.).

## Author contributions

Andres Nevarez, Conceptualization, Formal analysis, Investigation, Methodology, Funding acquisition, Writing - original draft, Writing - review and editing; Anusorn Mudla-Cell line creation; Sabrina Diaz cell culture and migration analysis; Nan Hao, Conceptualization, Resources, Formal analysis, Supervision, Funding acquisition, Investigation, Methodology, Project administration, Writing - review and editing.

