## Supplementary figures and images for "Using Deep Learning to Decipher the Impact of Telomerase Promoter Mutations on the Morpholome"

### Fig6B_PHATE_IFC_Both_TPM_rotating_plot.gif

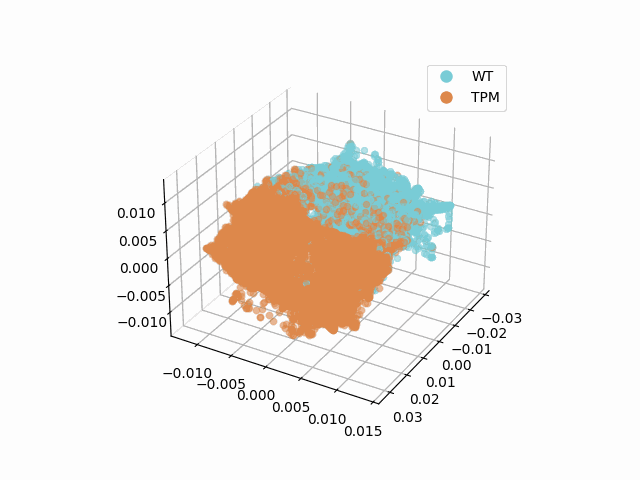

### Fig7B_IFC_Both_rotating_plot.gif

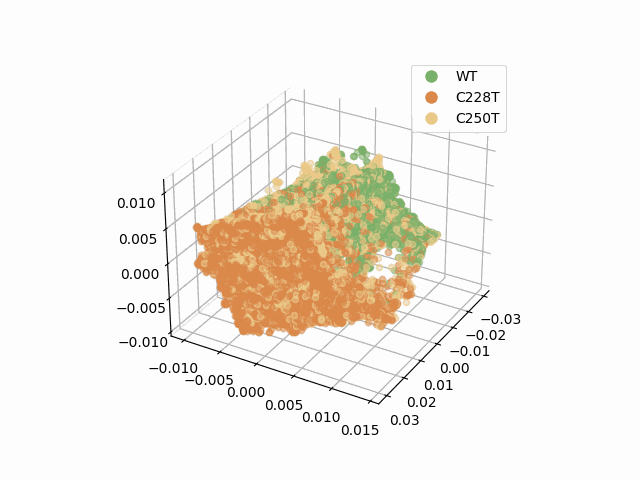

### SuppFigAB.pdf

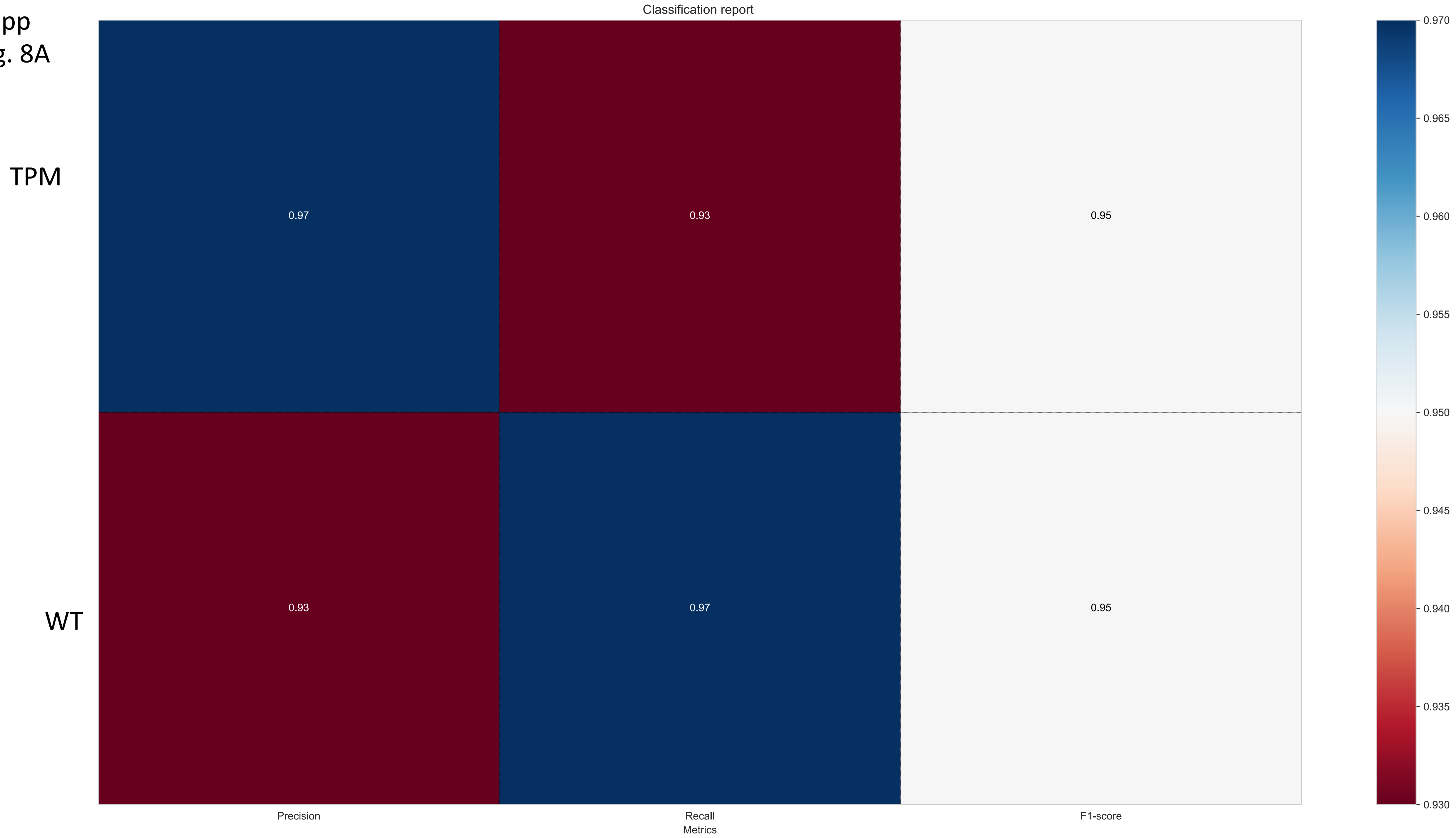

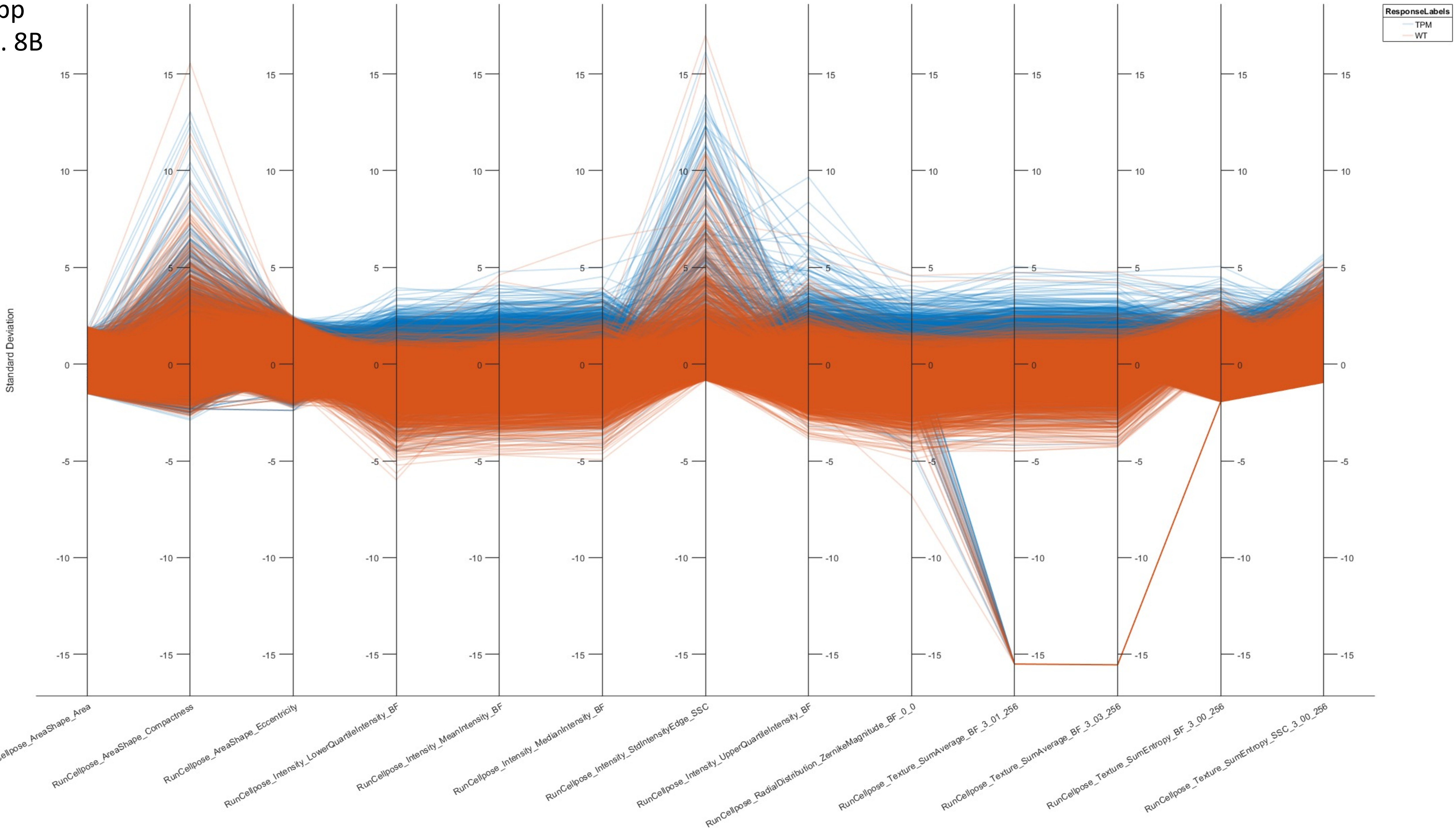

### SuppFigure3.pdf

#### Supplemental Figure 8

A

B

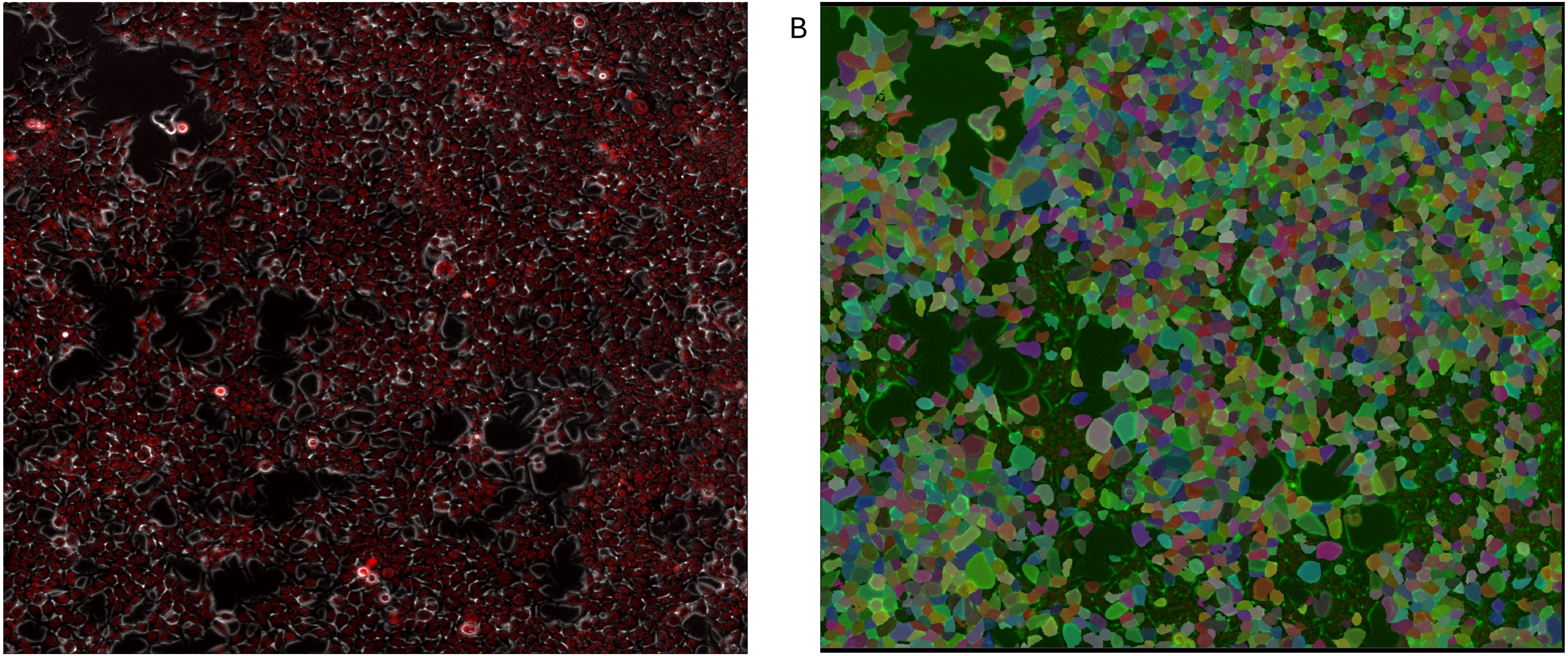
